## Supplemental Materials for "Using an Ordinary Differential Equation model to separate Rest and Task signals in fMRI"

### 1 Supplementary information

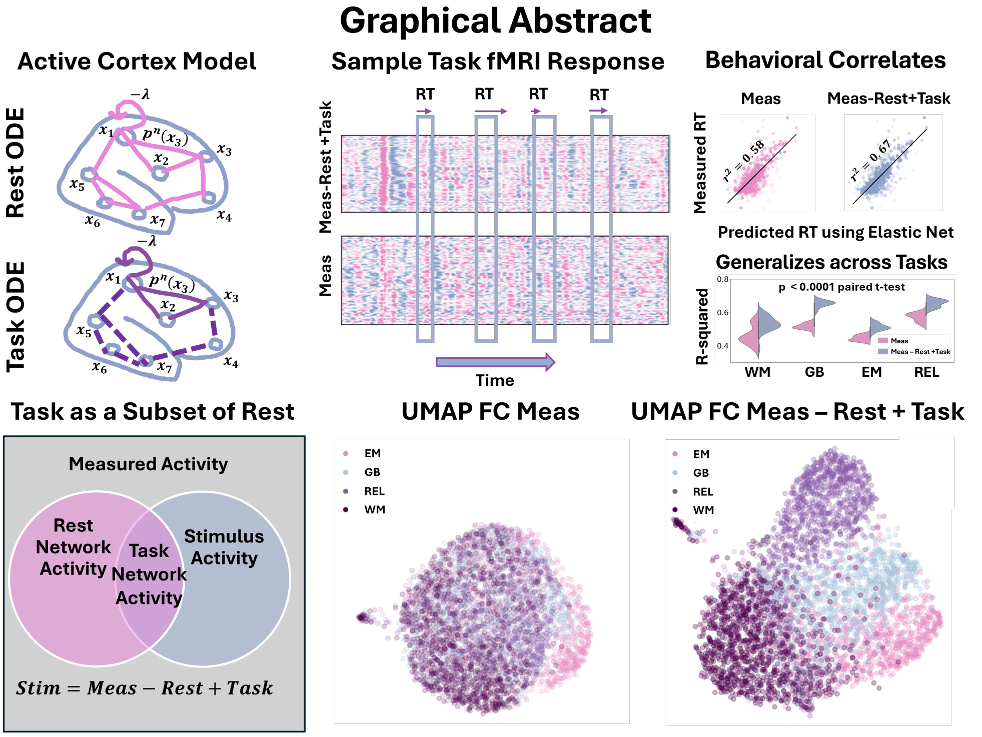


**Fig. 1** **Graphical Abstract** Top Left: The Rest and Task network ODEs. These network models are estimated from their respective fMRI data using data-driven methods that approximate Taylor series polynomials for each network edge, with the Task ODE forming a subset of the Rest ODE. Bottom Left: The Active Cortex Model (ACM) - The Stimulus Activity couples with the Resting State Networks to produce Task Network activity. To isolate the stimulus-related activity associated with the behavioral task, the Resting State Network activity modeled by its corresponding ODE is subtracted from the measured time series, while the Task Network activity, also represented by an ODE, is added back as shown in the Venn Diagram. Top Middle: A sample measured Task fMRI (unseparated) and Task isolated fMRI using the ACM (separated). Top Right: The reaction times for each trial have about 9 percent higher $R^2$ with the separated data vs the measured data on average across four HCP Tasks (p<0.0001). Bottom Right: By removing the Rest processes not present in the specific Task, the functional connectivity separates in high dimensional space providing more interpretability between distinct Tasks visualized using a 2D UMAP. Using an SVM, we quantify a 20 percent increase in accuracy in the UMAP space between the FC unseparated and FC separated. HCP Task abbreviations and the number of subjects used for Task separation: EM- Emotion (N=1017), GB- Gambling (N=1032), REL- Relational (N=1013), WM- Working Memory (N=1033)

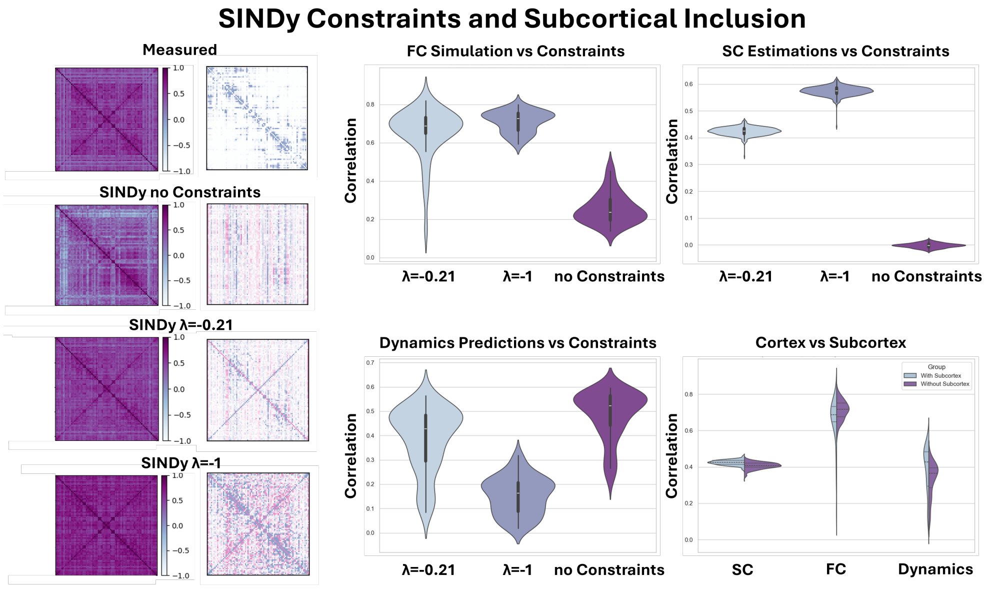


**Fig. 2** **SINDy Constraints and Subcortical Inclusion**: Comparison between different constraint options using the SINDy algorithm. Left panel, the simulated FC and 1st Order Coefficient are provided for no constraints, constraint close to the decay rate for the measured HRF, amd a large constraint of -1 (N=100 for SINDy Coefficients and N=90 simulation for averaged FC). No constraint has the best fit for the dynamics, but the coefficients have no similarity to the $SC$, whereas the larger constraints tend to lead to solutions where the coefficients are more similar to the $SC$ (N= 90 for Dynamics and N=443 for SC comparison). Close to the decay rate of the measured HRF we have a good fit for both the dynamical and $SC$ metrics. In the bottom right, the simulated FC, estimated $SC$ and predicted dynamics are compared for the inclusion of subcortical regions (N= same as for other panels respectively). The inclusion of the subcortex has a profound effect of improving the prediction of dynamics of other regions ($p < 0.00001$ two sided student t), while the other metrics do not change too much. Abbreviations: HRF- Hemodynamic Response Function. FC- Functional Connectivity. SC- Structural Connectivity. The violin box plot in C) shows the median as a white dot, the interquartile range as a thick bar from the 25th to 75th percentile, and whiskers extending to the most extreme data within 1.5 $\times$ IQR of the quartiles.

**Table 1** Brain Regions by Hemisphere in the order that they appear in FC and SC matrices

| No. Right | No. Left | Region |
|---|---|---|
| 1 | 84 | Cerebellum |
| 2 | 83 | Thalamus |
| 3 | 82 | Caudate |
| 4 | 81 | Putamen |
| 5 | 80 | Pallidum |
| 6 | 79 | Hippocampus |
| 7 | 78 | Amygdala |
| 8 | 77 | Accumbens |
| 9 | 76 | Insula |
| 10 | 75 | Entorhinal |
| 11 | 74 | Parahippocampal |
| 12 | 73 | Temporal Pole |
| 13 | 72 | Frontal Pole |
| 14 | 71 | Fusiform |
| 15 | 70 | Transverse Temporal |
| 16 | 69 | Lateral Occipital |
| 17 | 68 | Superior Parietal |
| 18 | 67 | Inferior Temporal |
| 19 | 66 | Inferior Parietal |
| 20 | 65 | Supramarginal |
| 21 | 64 | Bankssts |
| 22 | 63 | Middle Temporal |
| 23 | 62 | Superior Temporal |
| 24 | 61 | Postcentral |
| 25 | 60 | Precentral |
| 26 | 59 | Caudal Middle Frontal |
| 27 | 58 | Pars Opercularis |
| 28 | 57 | Pars Triangularis |
| 29 | 56 | Rostral Middle Frontal |
| 30 | 55 | Pars Orbitalis |
| 31 | 54 | Lateral Orbitofrontal |
| 32 | 53 | Caudal Anterior Cingulate |
| 33 | 52 | Rostral Anterior Cingulate |
| 34 | 51 | Superior Frontal |
| 35 | 50 | Medial Orbitofrontal |
| 36 | 49 | Lingual |
| 37 | 48 | Pericalcarine |
| 38 | 47 | Cuneus |
| 39 | 46 | Paracentral |
| 40 | 45 | Isthmus Cingulate |
| 41 | 44 | Precuneus |
| 42 | 43 | Posterior Cingulate |

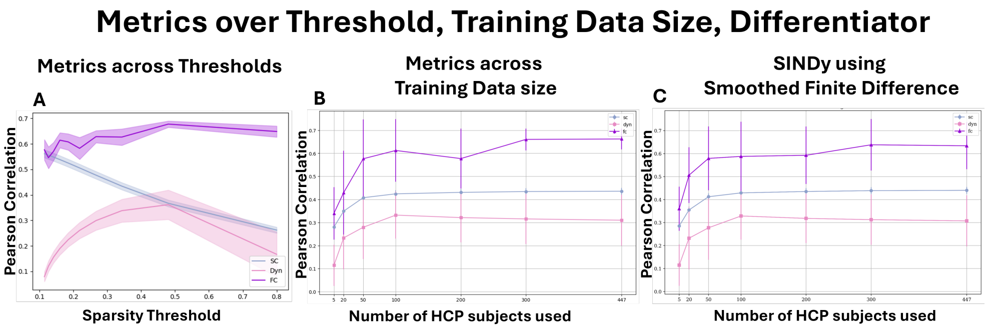


**Fig. 3 Metrics over Threshold, Training Data Size, Differentiator**: A) Looking at the performance of multiple metrics on the value of sparsity . While the correlation of the first-order coefficients with the $SC$, and the prediction of the derivative vastly change over the sparsity threshold range, the simulated functional connectivity barely changes across the whole range. Therefore, we are justified using $SC$ and dynamics as well as HRF as the main metrics to distinguish between models. B) The amount of training data needed for convergence. Around 100 HCP subjects with 4 sessions of Rest (1200 time points * 4 * 100) was the minimum required for convergence. C) Sample plot as center, but using the smoothed finite-difference differentiator instead of the spectral differentiator option in SINDy. The difference between them is not noticeable. The FC and Dyn were compared/ simulated to N=20 resting state scans. The SC were compared to N=443 DTI scans

.

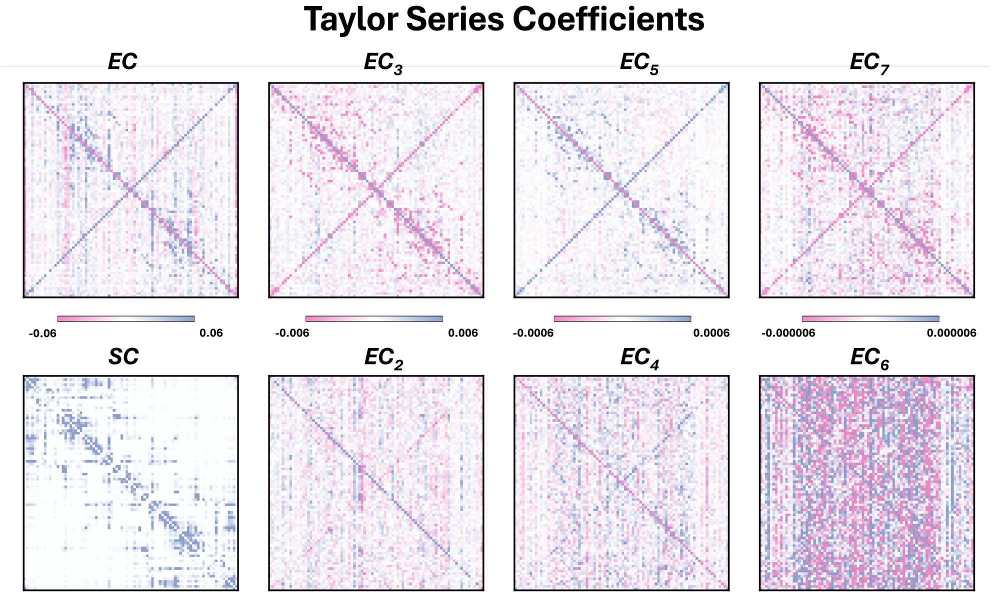


**Fig. 4 Taylor Series Coefficients:** A) The first seven Taylor series coefficients for $\lambda = -0.21$ computed by running SINDy on a 80 percent subset of resting state fMRI 447 individuals 100 times and averaging. Top and bottom have the same colorbar. The SC is shown bottom left in comparison. These are the complete coefficients shown in Figure 1 and at the bias-working point of the SINDy algorithm for fMRI. These coefficients were determined using 447 Resting State Subjects

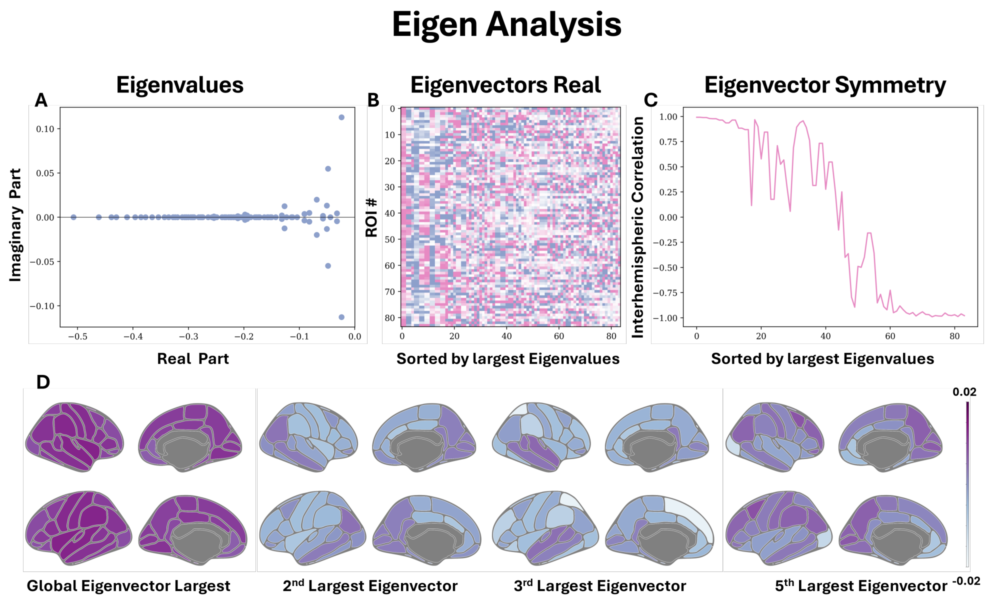


**Fig. 5** **Eigen Analysis**: A) The eigenvalues of the Jacobian (first order coefficients) shown on the top left are plotted on the complex plane. All eigenvalues have a real value less than one, suggesting the system is stable, although some modes have real components fairly close to 0 suggesting the system is close to criticality. B) The corresponding real component of the eigenvectors plotted column-wise in top middle, sorted by largest eigenvalue (closest to 0). C) The first eigenvectors show a global mode and are highly symmetrical across the hemispheres. The smaller eigenvectors shown in the top right are unimodal mode and are even anticorrelated with the contralateral component. D) In the bottom, the global eigenmode and some of the bilateral network eigenmodes are displayed for the entire cortex. The brain plots were made using ggseg

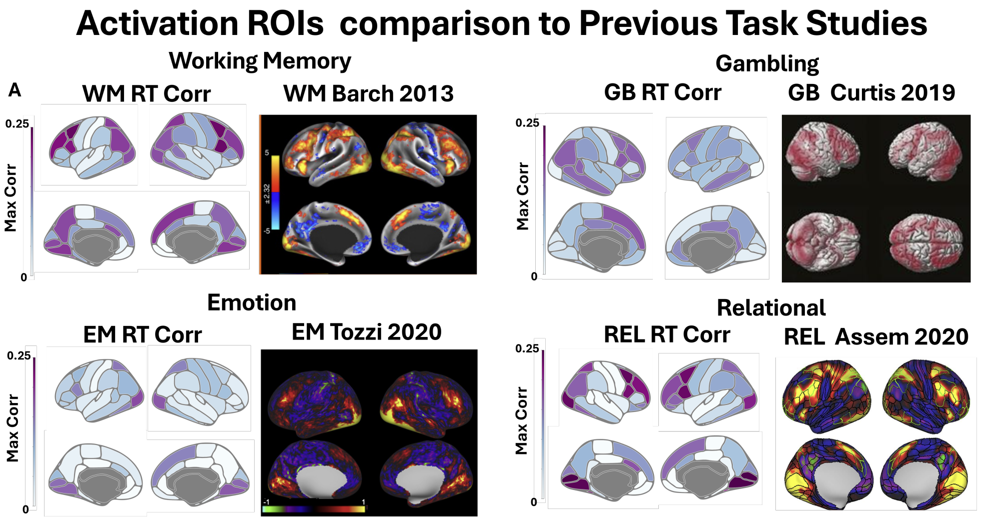


**Fig. 6 Comparisons to Previous Task Activation Maps:** Comparisons between our results spatially vs earlier GLM models on HCP Task data. Each of the previous Tasks Activation was determined by calculating GLM masks between two Tasks conditions or Task vs Baseline condition such as 0 vs 2 back for WM, or Fear vs Neut for EM. In Barch et al., 2013 the activation maps for Task vs Baseline and difficulty of Task were shown to be similar. The spatial maps shown from our study were calculated for the regions max correlation with the RT variable. These align well with Task conditions from previous work. Correlations are determined using N> 1000 scans for each Task. The brain plots were made using ggseg.

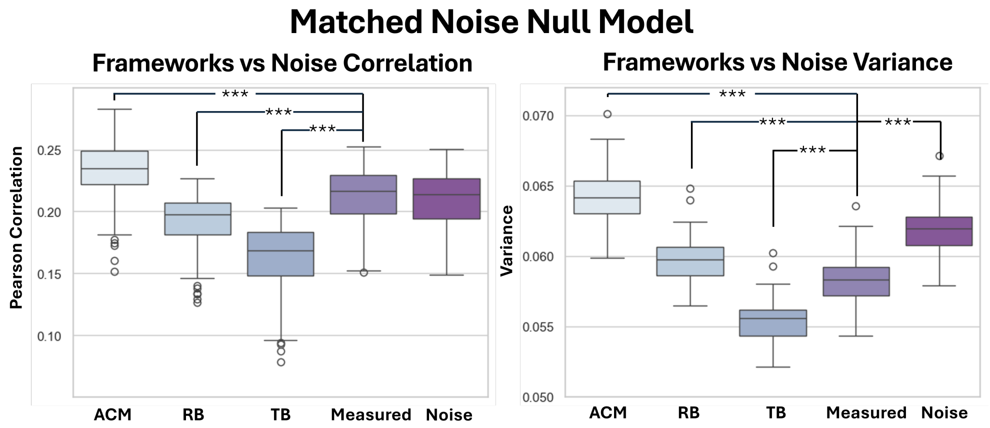


**Fig. 7 Matched Noise Null Model:** The Separation of the signal using various models. M. stands for Measured. The signal separation using both task and rest ODE, the Active Cortex Model (Rest - Task), is the only separation that yields significantly larger behavioral correlates (p $<$ 0.001) than the original Measured Signal. The matched signal represents subtracting the measured signal using a random signal with the same mean and variance as the Active Cortex Model. The figure on the left shows that the variance increases significantly ($p < 0.001$) for the Active Cortex Model after subtraction compared to the original and matched model. The p-values displayed are given with respect to the original measured signal, but are very significant $p < 0.001$ for all comparisons except for Measured vs Measured-Matched in the left plot and Measured vs Measured – Rest in the right plot. P-values computed using student t two sided test. N $>$ 1000 for Task correlation and p-value represents two sided paired t-test average 12 spatial temporal location correlation

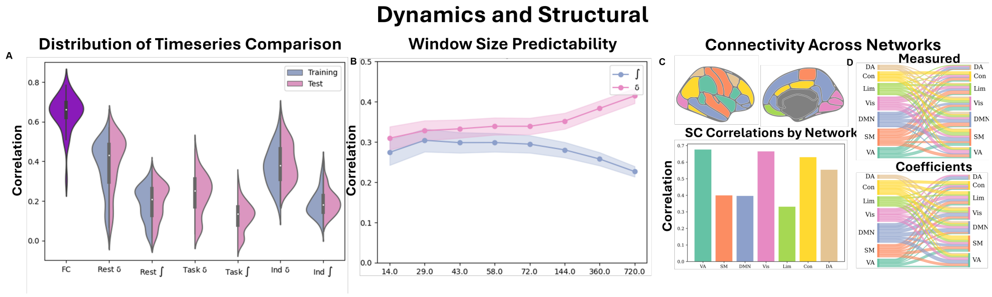


**Fig. 8 Dynamics and Strucutral Comparisons:** A) Comparisons between many different varieties of predicted timeseries and the respective measured ones from the fitted equation. Rest and Task refer to the dataset types averaged across individuals, while Ind refers to predictions to Rest averaged across ROIs. The correlation between simulated FC and measured FC in the far left. The unseen 90 resting state scans were used to test the data while a evenly matched 90 resting state scans were used to evaluated the training set. B) The average correlation computed over different time intervals for both the derivative and the original signal (denoted using the integral symbol) predicted by SINDy against the respective measured signals (averaged across ROIs). C) Top part: The seven Yeo cortical networks are modified to match Desikan Killiany parcellation (DMN - Default Mode Network, VA -Ventral Attention, DA - Dorsal Attention, SM- Somato Motor, Con - Control, Lim - Limbic, Vis - Visual). Bottom part: The correlation of each of the networks separately calculated from the estimated first-order and the $SC$ matrix. D) Sankey plots for the white matter $SC$ and the coefficients from the first-order Taylor expansion.

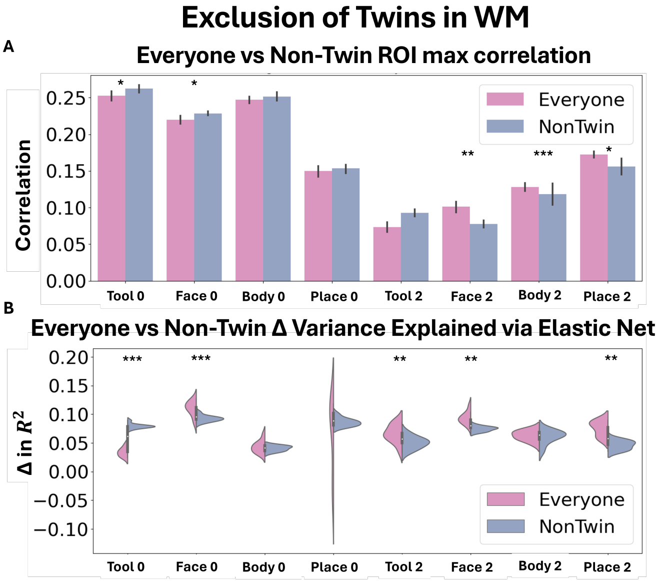


**Fig. 9 Twin Experiment:** A)Repeating the analysis of separating via ACM on a dataset that included everyone (N=1032), as shown in **??** or redoing the analysis on a set of non-twin HCP data (N=528) and testing on the twin subset (N=504). The y-axis is the correlation to RT and the x-axis is the various sub-tasks. While there are a few significant changes, they go both ways, and the results in general show that both analysis perform relatively similarly to each other. B) Observation of the change in variance according to the ACM model for the two datasets compared to the unseparated baseline model. Again, while some sub-tasks show significant differences, there is no overall trend. The dataset for the entire group showed a mean variance increase of 6.9 and the non-twin subset is 6.6 for WM. The stars have different significance values: *** $< 0.001$, ** $< 0.01$, * $< 0.05$ on a student t two sided test.

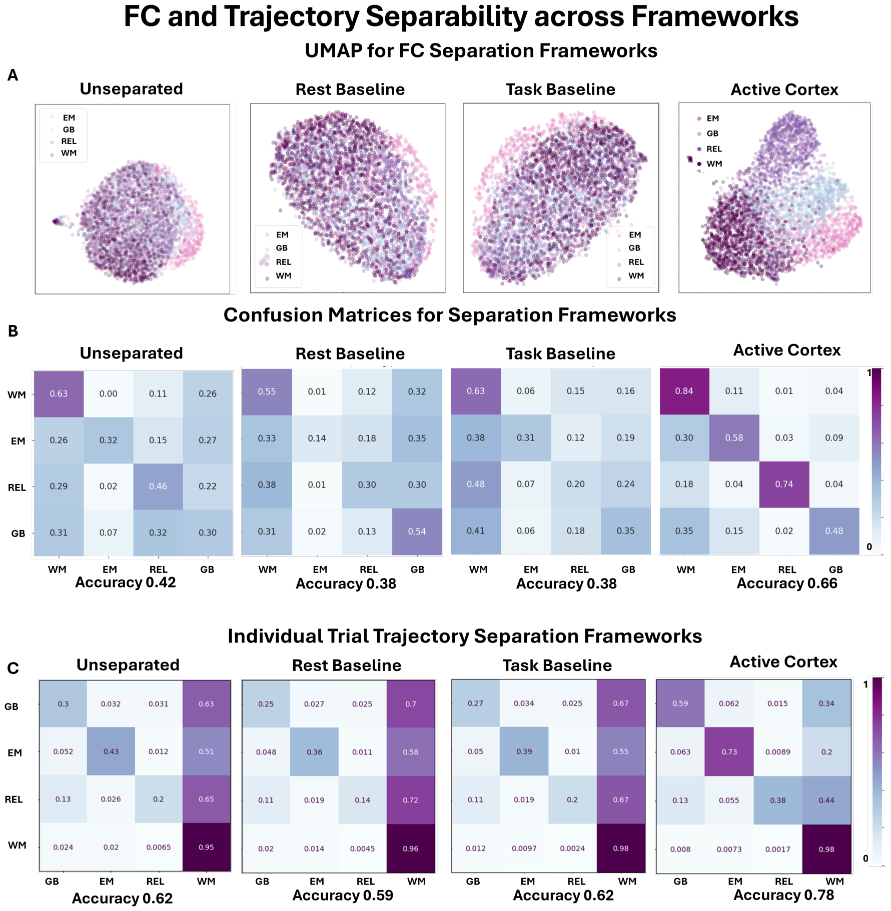


**Fig. 10 FC and Trajectories Separability across different Frameworks:** A) The FCs for the different frameworks of separability computed across all Task trials for a single individual (one point is one individual for the particular Task, N>1000). Only the Active Cortex model shows differentiation between the different Task FCs. B) The confusion matrices in the attempt to separate out the data in the UMAP embedding space using a SVM model. The other baseline models perform worse than the unseparated measured signal. The active cortex model shows a twenty percent increase in accuracy of the resulting SVM model over the unseparated cases. C) The separability was also tested using the individual trials from each category, rather than computing the FC across all individual trials from each category. A random forest classifier using 10 fold cross validation was used to test how well they were separated for each type of separation framework. The ACM once again is the only tested framework that improves from the baseline accuracy by 15 percent while other frameworks show no benefit. For all the data the N for each Task was greater than 1000.

.

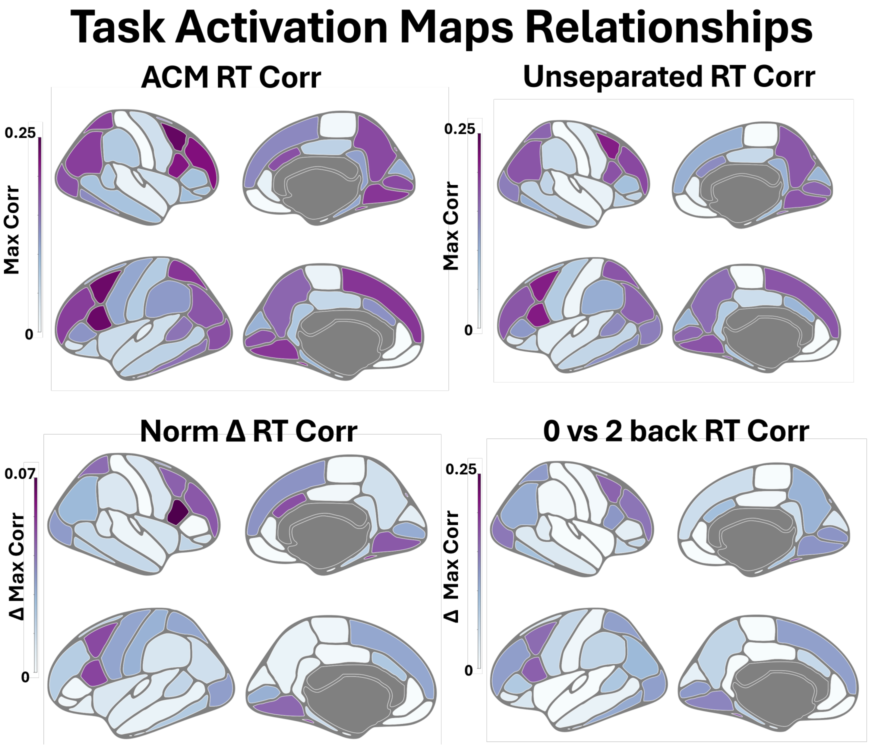


**Fig. 11** **Task Activation Maps Relationships:** Top Left: The activation Map from ACM based on the max correlation of a ROI across time for 0 back WM (N=1032). Top Right: The activation Map from the unseparated data based on the max correlation of a ROI across time for 0 back WM (N=1032). Bottom Left: The normalized change in correlation between the ACM (separated) and the unseparated cases showing a subset of regions that were originally identified in the Task increasing in correlation (N=1032). Bottom Right: The difference between the activation maps based on correlation to RT for the zeroback and twoback (N=1032). The zeroback trials are more correlated with the RT, and the difference between them removes associations between regions that are necessary for the Task such as the visual cortex but do not reflect the Task severity. The change in separated vs unseparated cases match the difference between tasks suggesting that the increases in correlation are in regions that are specific to the particular difficulty of the Task, not just regions that activate for Task completion such as the visual cortex to see the input or the motor cortex for button pushing. Significance Test: Norm $\Delta$RT Corr to 0 vs 2back = 0.79 $R^2$ Norm $\Delta$RT Corr to ACM RT Corr = 0.75, Norm $\Delta$RT Corr to Unseparted RT Corr = 0.76. P-value difference between 0.79 $>$ 0.75 is 0.002 and 0.79 $>$ 0.76 is 0.003.

.

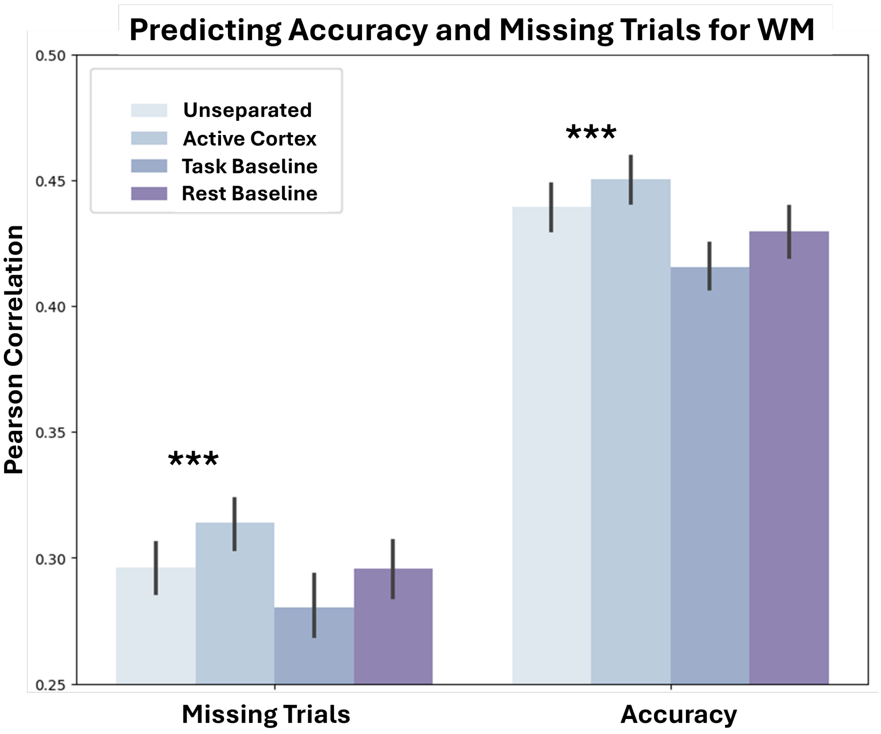


**Fig. 12 Predicting Missing Trials and Accuracy for WM trials**: The percentage of missing trials and accuracy was calculated for each individual across both WM runs. Functional connectivity (FC) was computed for all the different separation strategies, as visualized using the UMAP in Figure 6. The FCs were then mapped to the percentage of missing trials and accuracy using an Elastic Net. The results of a 100-fold cross-validation scheme are shown above for each framework strategy using a 80/20 split (N=100 for each bar). The ACM performed significantly better at predicting both missing trials and accuracy, although the difference is smaller in magnitude compared to the differences observed in RT. The stars have different significance values: *** $< 0.001$, ** $< 0.01$, * $<$ 0.05 on a student t two sided test

.

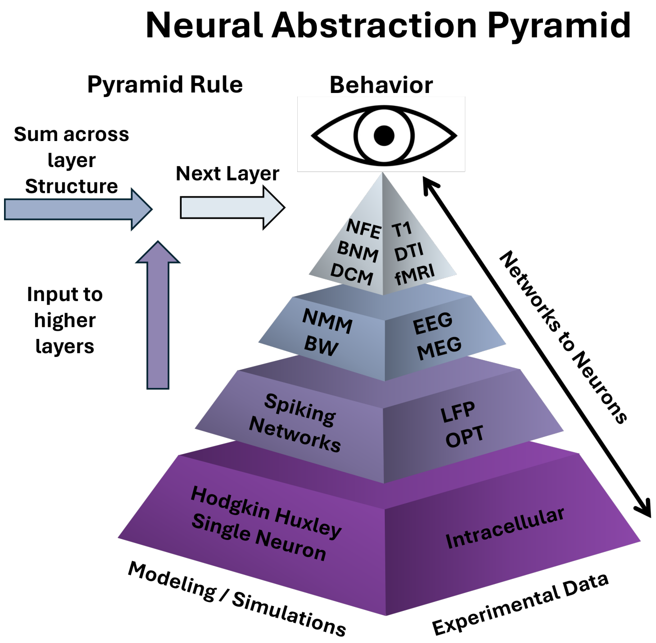


**Fig. 13** The Neural Abstraction Pyramid representing different layers of neural activity from neurons to the whole network. Behavior is represented at the top most layer as it emerges from the activity of different components of the pyramid. Legend: BNM -Brain Network Model, NFE- Neural Field Equations, fMRI - functional Magnetic Resonance Imaging, DCM - Dynamical Causal Modeling, T1 - Structural Image, DTI - Diffusion Tensor Imaging, NMM -Neural Mass Models, EEG-Electroencephalogram, MEG - Magnetoencephalography, BW - Balloon Windkessel Model, LFP - Local Field Potential, OPT- Optogenetics Image Ref
